# Goitered Gazelle’s (*Gazella Subgutturosa*) Habitat Desirability Modeling by Using Maximum Entropy (Maxent) Method

**DOI:** 10.1101/2022.02.10.479956

**Authors:** Abbas Naqibzadeh, Jalil Sarhangzadeh, Ahad Sotoudeh, Marjan Mashkour, Judith Thomalsky

## Abstract

The models predicting the spatial distribution of species can simulate the suitability of species habitats on different spatial scales, based on species records and site characteristics to gain insight into ecological or evolutionary drivers or to help predict habitat suitability across large scales. Species distribution models (SDMs) based on presence-absence or presence-only data use widely in biogeography to characterize the ecological niche of species and to predict the geographical distribution of their habitat. Although presence-absence data is generally of higher quality, it is also less common than presence-only data because it requires more rigorous planning to visit a set of pre-determined sites. Among the algorithms available, one of the most widely used methods of developing SDMs is the Maximum Entropy (MaxEnt) method. The MaxEnt uses entropy to generalize specific observations of presence-only data and does not require or even incorporate points where the species is absent within the theoretical framework. The purpose of this study is to predict the suitable habitat for Goitered gazelle (Gazella subgutturosa) in the Samelghan plain in northeastern Iran. The results showed that the variables of the Mediterranean climate classes, slope 0-5% class and semi-dense pastures with type Acantholimon-Astragalus are more important than other environmental variables used in modeling. The area under curve (AUC), Receiver Operating Characteristic (ROC), and the classification threshold shows model performance. Based on the ROC (AUC=0.99) results in this study, it was found that Maxent’s performance was very good. Desirability habitat was classified based on the threshold value (0.0277) and the ROC, which approx 11% of the area, predicted suitable habitat for Goitered gazelle.

## Introduction

Habitats’ study has key importance for the development of wildlife conservation policies, evaluation, and conservation (Suleman *et al*., 2020). Ecological studies such as; Habitat Suitability models provide plenty of knowledge about the relationships of wildlife with habitats (Allouche *et al*., 2008; Kuloba *et al*., 2015). Species distribution models (SDMs) are increasingly being used in environmental management, such as habitat suitability studies (Luo *et al*., 2020), managing endangered species, and environmental change impacts (Hirzel *et al*., 2006). The Models predicting the spatial distribution of species (Guisan & Zimmermann, 2000; Pearce and Boyce, 2006; Ghanbarian *et al*., 2019) can simulate the suitability of species habitats on different spatial scales, based on species records and site characteristics (Kuloba *et al*., 2015; Qin *et al*., 2017; Mousazade *et al*., 2019) to gain insight into ecological or evolutionary drivers or to help predict habitat suitability across large scales (Elith & Leathwick, 2009; Kramer-Schadt *et al*., 2013). Therefore, modeling species distribution has become a key tool in conservation (Allouche *et al*., 2008; Kuloba *et al*., 2015), ecology (Phillips & Dudík, 2008) evolutionary biology (Guisan & Thuiller 2005; Kozak *et al*., 2008; Dormann *et al*. 2012; Muscarella *et al*., 2014), biogeography (Elith *et al*., 2011), evolution, invasive species control (Phillips *et al*., 2006) and wildlife management studies (Long *et al*., 2008; Franklin, 2013; Wan et al., 2016; Zhang et al., 2019a; Li et al., 2020).

In recent years, many techniques for modeling species’ niches and distributions have been developed (Guisan & Zimmermann, 2000; Guisan & Thuiller, 2005; Kozak *et al*., 2008; Peterson *et al*., 2011; Radosavljevic & Anderson, 2014). Computer tools such as a large number of algorithms and software platforms (Guilbault *et al*., 2019) with Geographic Information System (GIS) (Traill & Bigalke, 2007), and statistical modeling techniques to draw up predictive maps (Jiménez-Valverde *et al*., 2007; Naqibzadeh et al,. 2021) have been become more widely used in ecology and conservation biology (Guisan & Zimmermann 2000; Warren *et al*., 2008; Kamyo & Asanok, 2020). One of the main shortcomings of distribution model predictions is a lack of reliable information on species absence (Jiménez-Valverde *et al*., 2007). Species distribution models (SDMs) based on presence-absence or presence-only data (Guilbault *et al*., 2019) are used widely in biogeography to characterize the ecological niche of species and to predict the geographical distribution of their habitat (Naimi et *al*., 2014). The presence-only data only contains information about species presence, in contrast to presence-absence data which records both where species have been found present and where they have not been found (Warton & Shepherd, 2010; Renner *et al*., 2015). Although presence-absence data is generally of higher quality, it is also less common than presence-only data because it requires more rigorous planning to visit a set of predetermined sites (Van Strien *et al*., 2013; Ruete & Leynaud, 2015; Guilbault *et al*., 2019). The presence-only data allow for easy public and private involvement in biological monitoring and are the dominant source of species occurrence data (Elith *et al*., 2011; Pédarros *et al*., 2020).

Among the algorithms available, one of the most widely used methods of developing SDMs is the Maximum entropy (MaxEnt) method (Phillips *et al*., 2017; Tourne *et al*., 2019). MaxEnt uses entropy to generalize specific observations of presence-only data and does not require or even incorporate points where the species is absent within the theoretical framework (Kamyo & Asanok, 2020). Maxent model is a useful technique to predict the potentially suitable habitat (Evcin *et al*., 2019; Qin *et al*., 2020), geographical species distribution (Phillips *et al*., 2006; Jiménez-Valverde, 2012; Merow *et al*., 2014; Xu *et al*., 2015; Mi *et al*., 2016; Fronczak *et al*., 2017; Wan *et al*., 2019; Wang *et al*., 2019; Kamyo & Asanok, 2020) on the basis of the most significant environmental conditions (Phillips *et al*., 2006; Moreno *et al*., 2011; Tourne *et al*., 2019). MaxEnt model on simulating the suitable geographical distribution of species, has more advantages than other models, including a good performance with incomplete datasets, short model running time, easy operation, small sample size requirements, and high simulation precision (Hernandez *et al*., 2006; Phillips *et al*., 2006; Pearson *et al*., 2007; Ortega-Huerta and Peterson, 2008; Li *et al*., 2020).

Many habitats outside the area under the management of the Department of Environment are of particular importance, especially in providing shelter and food for wildlife, which can provide a good place for fauna and flora. The purpose of this study is to model the habitat of Goitered gazelle in these areas with the MaxEnt method. It should be noted that many of these habitats are important in wildlife conservation and management, and should be considered in conservation discussions of fauna and flora. The importance of this region is due to the existence of the Rivi archaeological site in the basin because based on the samples obtained from zooarcheological studies, the historical distribution of species can be understood. Also, with zooarchaeology and analysis of animal remains, it is possible to reconstruct past habitats (Mashkour *et al*., 1999, Mashkour, 2001; Mashkour, 2013; Neer, 2017). Indeed, this study may serve as a pilot project and will give a basic outline for future investigations along with the Tappe Rivi Project that will estimate wildlife versus animal management through ancient times and in comparison to modern fauna.

## Matrial and Methods

Modeling can be made by using many variables in wildlife studies. MaxEnt model can use environmental variables and presence-only data (Evcin *et al*., 2019) to calculate the constraints and explores the possible distribution of maximum entropy under this condition, and then predicts the habitat suitability of the species at the study area (Phillips *et al*., 2006; Merow *et al*., 2013; Zhang *et al*., 2019b).

The Samelghan plain (10 km south of Atrak valley) is located in Northern Khorasan province, North East of Iran, and exhibit an ancient settlement area of more than 100 ha with four major Tappe ruins. The archaeological remains date back from the Late Bronze and Iron Age throughout the early historic periods of the Achaemenid and sasanid empire (approx. 1500 BC until 500 AD), plus a succeeding village-like occupation during Early Islamic times (Jafari & Thomalsky 2016; Thomalsky, 2016). The study area covers an area of 111660 hectares (Figure 1), which is located in the geographical position of 37° 21’ to 37° 40’ north latitude and 56° 26’ to 57° 06’ east longitude.

**Fig1.**
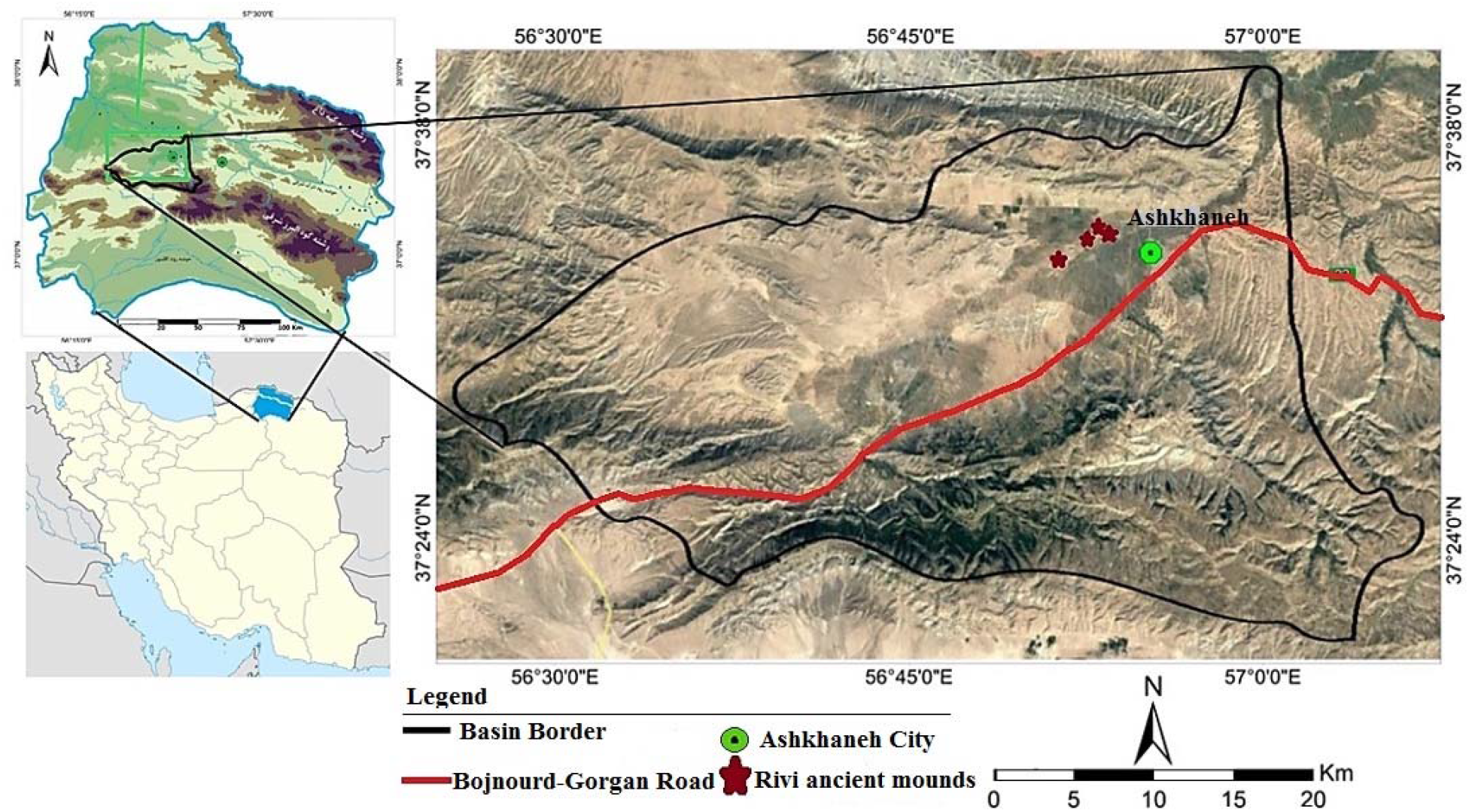
the location of the study area in Iran

The environmental (classified) variables used for modeling include: topography and geomorphology, climatology, land-use, vegetation, water resources, and human development variables such as villages and roads. Also, class maps of slope percentages and the main aspects were prepared by using DEM. All variables were converted to raster maps after digitization with 30×30 m cell size. In order to collect occurrence records, the distance sampling method (Waltert *et al*., 2008; Thomas *et al*., 2010), direct observations, footprints, repose imprints, feces, and tracks were used. One of the advantages of the MaxEnt as a modeling method is that it is able to work even with a very small sample size (Pearson *et al*., 2006; Phillips *et al*., 2006; Phillips & Dudík, 2008; Saupe *et al*., 2015; Zhang *et al*., 2019b; Karami *et al*., 2020). In this research, the geographical coordinates of the point were recorded using the Global Positioning System (GPS) as the presence point, and a total of 43 points were obtained for the species of Goitered gazelle in the Samelghan plain. After closing the border of the region based on the watershed, all variables were prepared according to the border.

## Results and Discussion

MaxEnt model produces this data: Prediction Maps (Prediction Maps); response curve (Response Curves) with an AUC (Area Under Curve); and Jackknife’s results of analysis help researchers to interpret and understand the outcomes of the MaxEnt model.

MaxEnt model output is logistic, which assigned each grid cell of the study area values ranging from 0 (completely unsuitable) to 1 (fully suitable) (Signorini *et al*., 2014; Alcala-Canto *et al*., 2018; Huercha *et al*., 2020). therefore, that if this value reaches 1, the habitat has higher desirability for the species and, conversely, reaches zero, the habitat desirability is reduced (Gormley *et al*., 2011) (Figure 2).

**Fig2:**
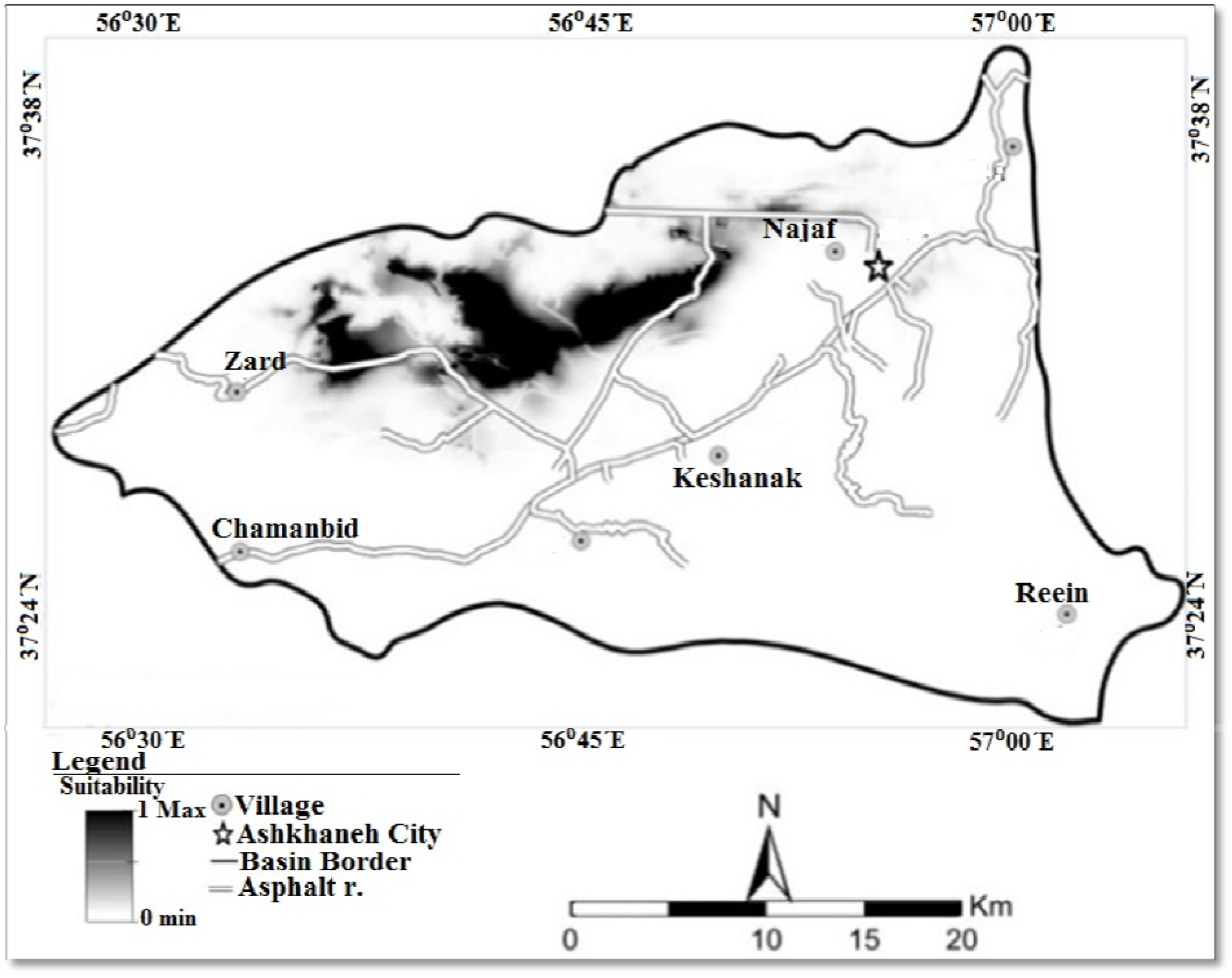
The Habitat continuous map

### Jackknife plot and response curves

MaxEnt can also estimate occupancy (referred to as ‘logistic output’) if the above assumptions are met, and users have additional knowledge of the occurrence probability of a species under ‘average’ conditions (Phillips & Dudík, 2008; Yackulic *et al*., 2013).

To evaluate the accuracy of each model, we used the threshold-in-dependent Area Under the Curve (“AUC”) calculated from the “Receiver Operating Curve” (ROC) (Fielding & Bell, 1997; Pearce & Ferrier, 2000). The ROC curve plots sensitivity against 1-specificity for all possible values of the threshold habitat suitability above which the habitat is assumed to be suitable (Fielding & Bell, 1997). The AUC can be interpreted as the probability that a randomly chosen presence site will be more highly ranked than a randomly chosen absence one (Pearce & Ferrier, 2000; Merow *et al*., 2013; Alatawi *et al*., 2020). An area under the receiver operating characteristic curve was examined for additional precision analyses (Evcin *et al*., 2019). The area under the curve (AUC) of the receiver operating characteristics (ROC) is a recommended index for model validation (Fielding & Bell, 1997; Moreno *et al*., 2011; Merow *et al*., 2013; Fand *et al*., 2014; Ghanbarian *et* al., 2019; Huercha et al., 2020) as well as is a common way to assess the performance of a species distribution model with binary response (Pédarros *et al*., 2020).

Briefly, The ROC plot ranges from 0.5 to 1.0. Values close to 1 means better performance (Phillips et al. 2006; Yuan *et al*., 2015; Ghanbarian *et* al., 2019; Wan *et al*., 2019; Huercha *et al*., 2020; Kamyo & Asanok, 2020). Values of ROC are described as an excellent model between 0.9 and 1, good between 0.8 and 0.9, fair between 0.7 and 0.8, and poor below 0.7 (Alcala-Canto et al. 2018; Preau *et al*., 2018; Evcin *et al*., 2019; Huercha *et al*., 2020; Ito *et al*., 2020). In this model, the area below the curve for the Goitered gazelle species was 0.990, with a standard deviation of 0.001, which indicates the ability to detect very good performance. (Figure 3).

**Fig3:**
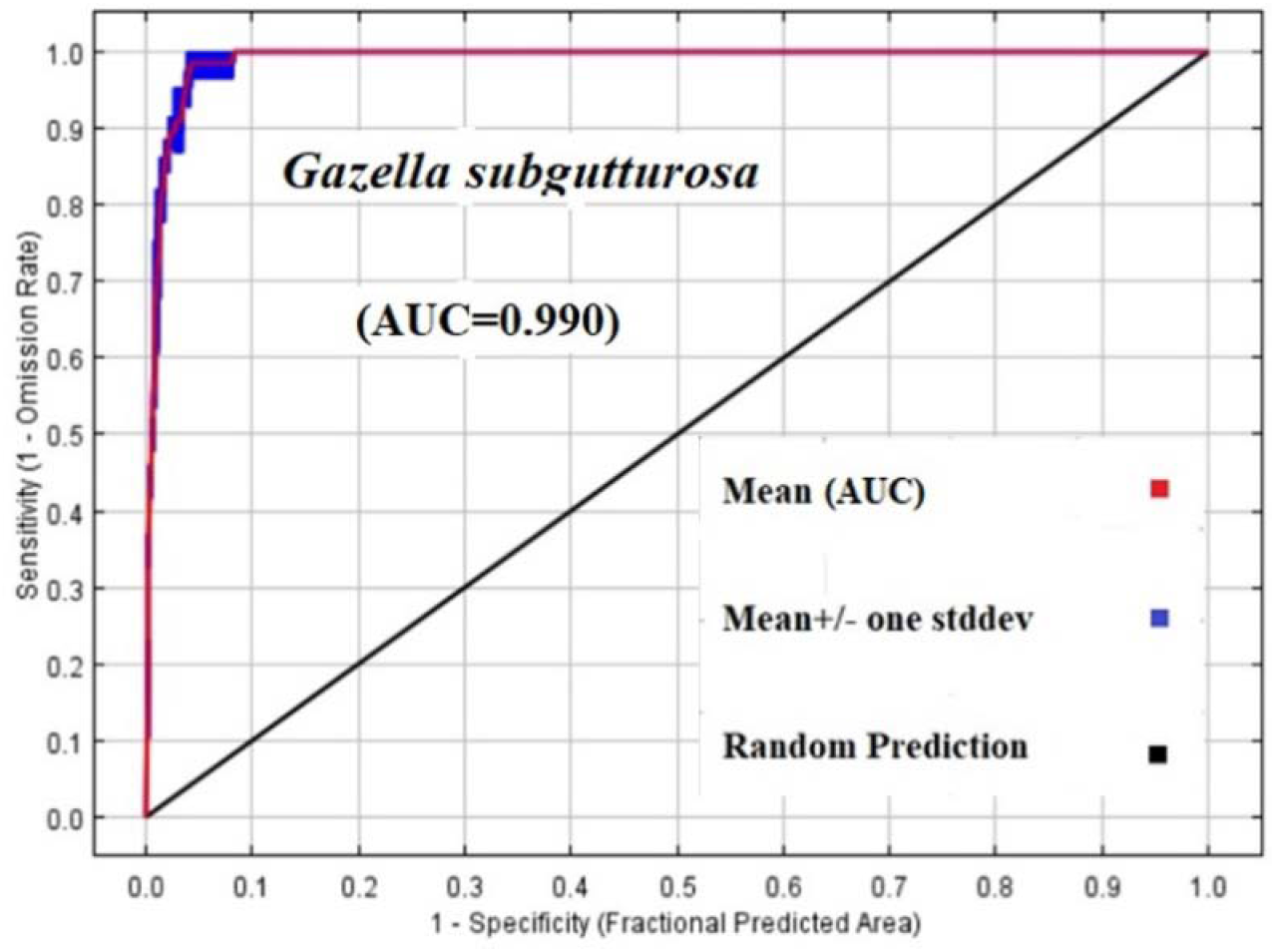
the ROC curve and AUC value

The jackknife plot validation model was used for the validation of model (Pearson *et al*., 2007) The jackknife plot was used to assess the relative importance of the variables (Phillips *et al*., 2006; Zhang *et al*., 2018; Evcin *et al*., 2019). The assessment capacity of species distribution model by AUC is criticized because of its relativity (Lobo *et al*., 2008). The AUC may be a good statistical measure of discrimination ability but it often fails to quantify the ecological likelihood of modelled distribution especially when estimated from presence-only data (Lobo et al., 2008; Pédarros *et al*., 2020). The Jackknife plot shows the importance of variables in three different colors; The blue color indicates how much of the species information is justified when running the model with only one variable, the light green indicates the implementation of the model without the desired variable, and the red color indicates the implementation of the model with all variables (Phillips *et al*., 2004). The Jackknife plot generally shows how environmentally friendly variables are effective in species distribution, based on The Jackknife plot, which can be used to determine which variable alone is most influential in modeling. Thus, the distance from the Mediterranean climate is the most effective variable in the development of Goitered gazelle species modeling (Figure 4) which can be removed by The greatest decrease occurs in the amount of AUC.

**Fig4:**
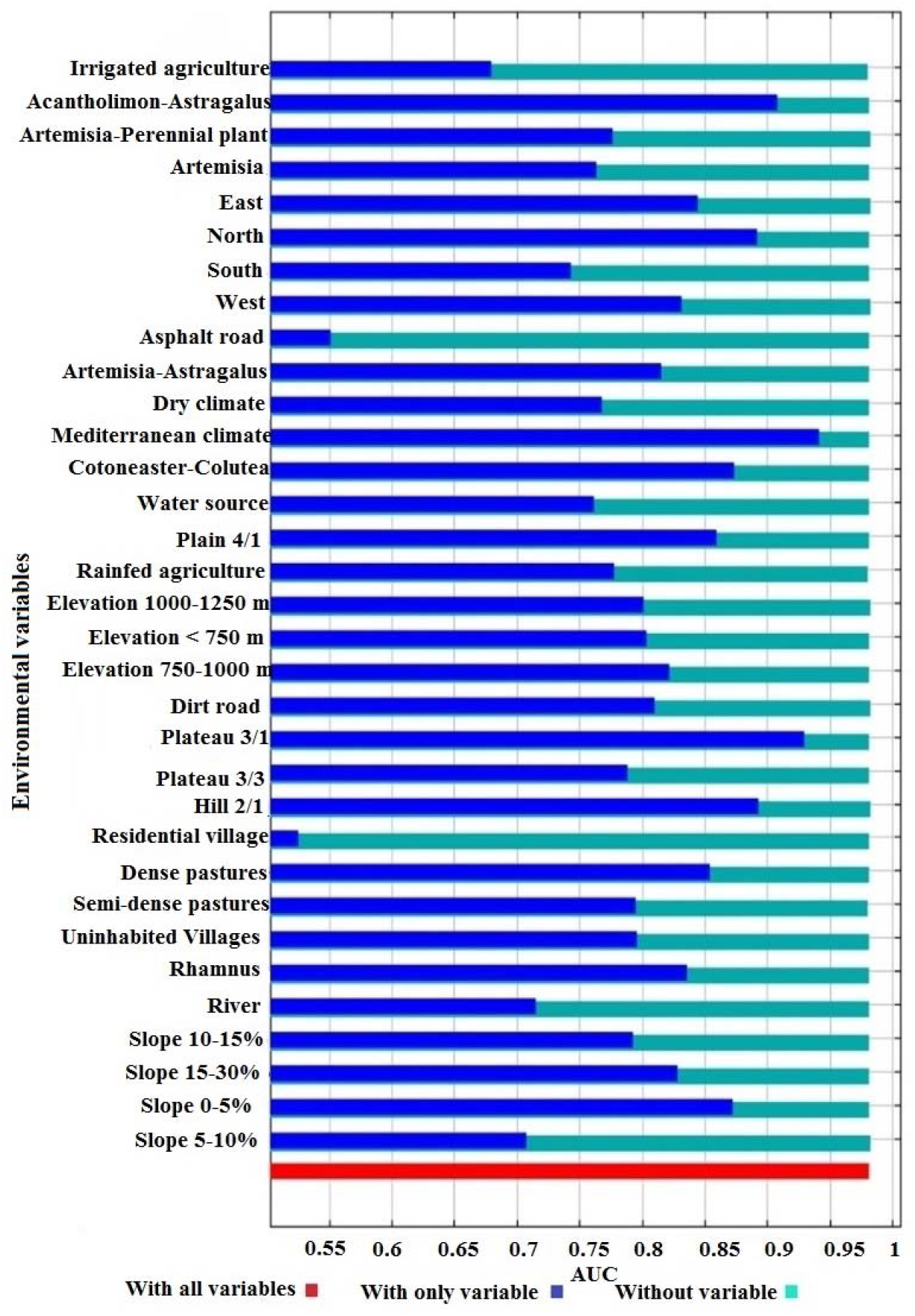
the jackknife test in determining the importance of variables

If all variables are in their mean values, the response curve to each variable will be in the form of graphs in which the red color indicates the average value of each variable in the two repetitions and the blue color indicates the standard deviation. In the graphs related to the environmental variables, the X-axis indicates the distance from the desired variable and the Y-axis indicates the probability of species presence, which ultimately determines the habitat suitability for the species being studied. Figure 5, shows the Goitered gazelle response curve to environmental variables. According to Figure 5, environmental variables have different effects on the distribution and presence of the species and ultimately the suitability of the Goitered gazelle habitat. According to the response curves, the distance from the conditions of the Mediterranean climate variable increases the probability of the presence of Goitered gazelle and then has a linear effect on the probability of presence. The response curve to the distance from the slope of 5-0%, so that the more the distance from the environmental conditions with this feature (slope of 5-0%) is reduced, the probability of the presence of the species is reduced and as a result, the desirability of the habitat will be reduced. The Goitered gazelle ’s response to the semi-dense rangeland variable is similar to the 5-0% slope variable. However, as inferred from Figure 5, a slope of 15-30% is one of the factors that will increase the likelihood of the presence of the species by increasing the distance from it. Other variables also have an effect on the probability of species presence, which can be interpreted by responding to the positive or negative effects of habitat suitability.

**Figure 5:**
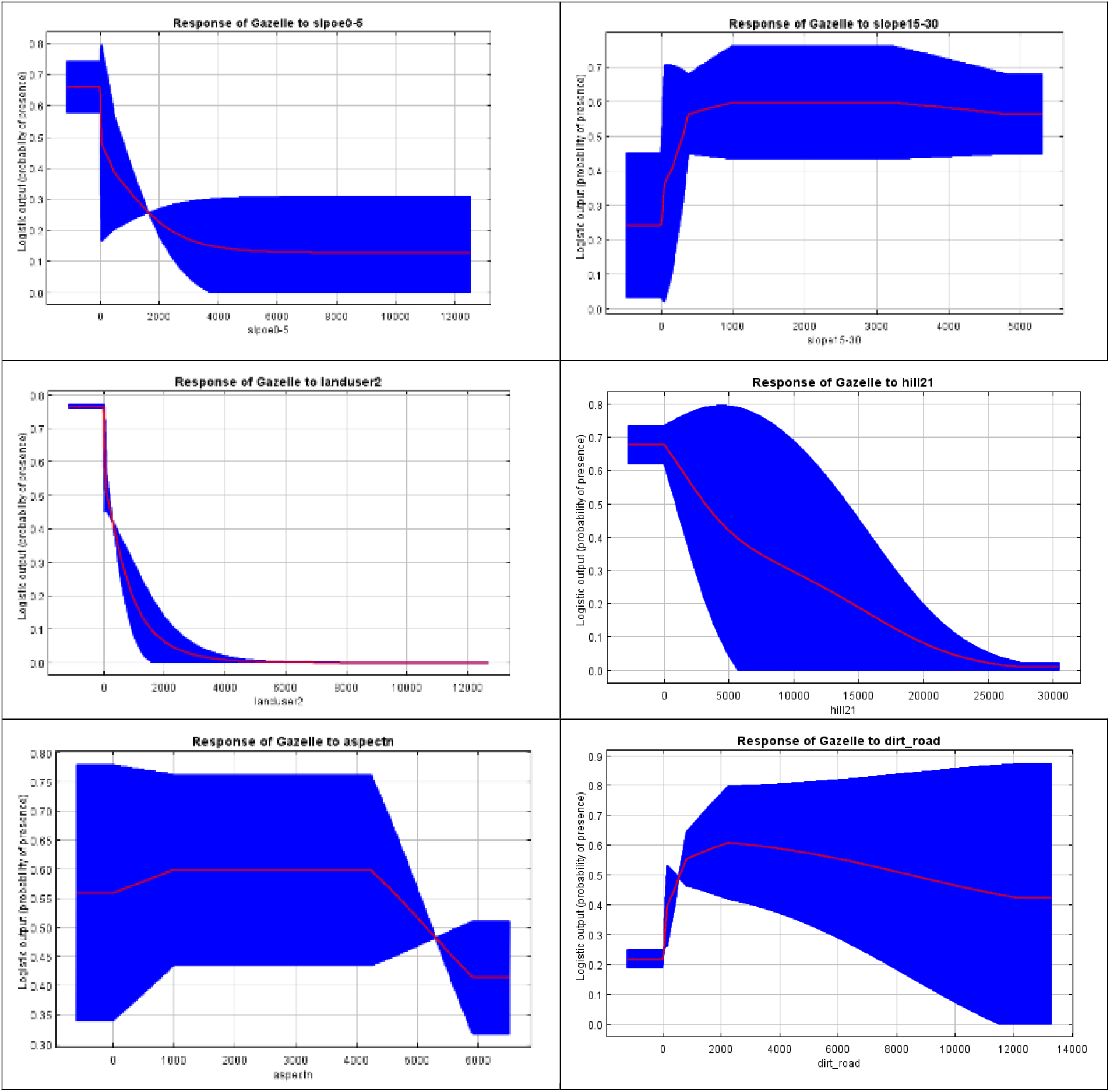
the response curves of Goitered gazelle to environmental variables

To determine the importance of each of the available variables from the statistics calculated by the software based on the share of each variable in the model (Phillips *et al*., 2012), the response curves of the variables for the single-variable model and the Jacqueline test for Evaluating AUC changes was used when deleting any variable (Yost et al., 2008). Table 1, show the relative share of each of the environmental variables in the distribution of Goitered gazelle at the surface of the Ashkhaneh watershed. As it is known, the distance from the slope of 5-0% and the distance from the semi-dense rangeland have the highest participation in Goitered gazelle distribution.

**Table 1:** Percent contribution values of variables used in modeling

| Variable | Percent contribution | Variable | Percent contribution |
| --- | --- | --- | --- |
| The Slope 0-5% | 34.3 | The slope 5-10% | 0.4 |
| Semi-dense pastures | 25 | Artemisia-Perennial plant | 0.3 |
| Dirt road | 6.1 | Southern Aspects | 0.3 |
| Western Aspects | 4.7 | Plateau 3/3 | 0.2 |
| Hill 2/1 | 4.6 | Western Aspects | 0.2 |
| The slope 15-30% | 4.4 | Eastern Aspects | 0.2 |
| Mediterranean climate | 3.7 | Asphalt road | 0.2 |
| Rainfed agriculture | 3.7 | Residential village | 0.1 |
| Plain 4/1 | 2.5 | The slope 10-15% | 0.1 |
| Northern Aspects | 2.5 | Uninhabited Villages | 0.1 |
| Irrigated agriculture | 2.2 | Dense pastures | 0.1 |
| Artemisia | 1.9 | Plateau 3/1 | 0.1 |
| Elevation <750 m | 1 | Elevation 1000-1250 m | 0.1 |
| River | 0.6 | - | - |

Maps for Goitered gazelle, was entered into the ArcGIS10.3 software by the Maxent software in ASC format and converted to a raster format so that according to the threshold value obtained from the model 0.0277 for Goitered gazelle to be classified into two desirable and undesirable classes (Figure 6). High threshold values indicate optimal habitat and lower values than threshold indicate adverse habitat for both species.

**Figure 6:**
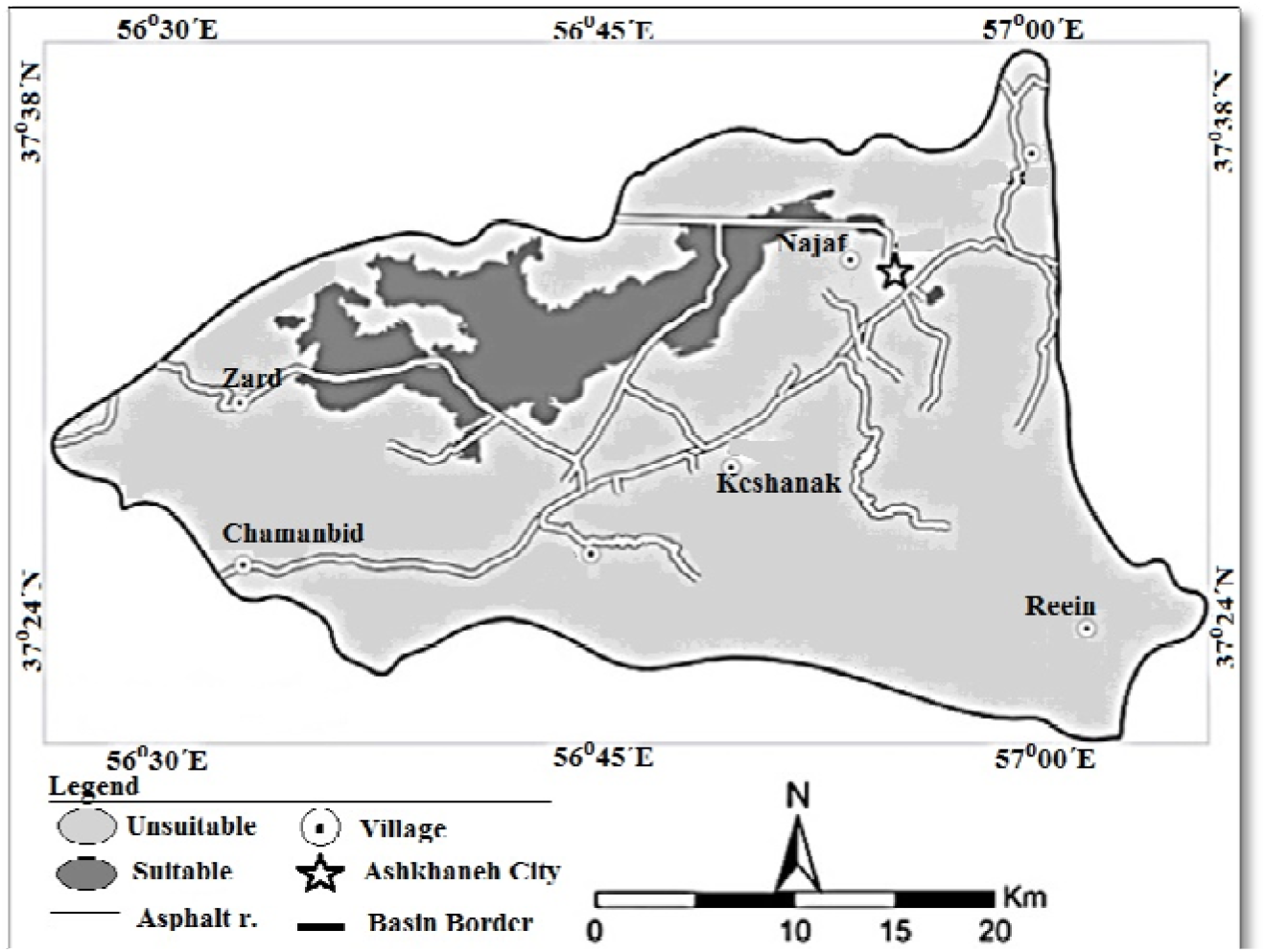
The Habitat classification map

## Discussion

The models provide a quantitative assessment of species extinction risk, identify potential threats to all animal species, and provide maps of the world’s mammal distribution points. Habitat evaluation and suitability provide strategic priorities for the protection of global mammals (Crooks *et al*., 2017).

Modeling methods can be used for different purposes, including; Determining the suitability of species habitats (Toor *et al*., 2011) Predicting the trend of species expansion at the level of a region (Giovanelli et al., 2010) as well as predicting high-risk areas of conflict between wildlife and human species (Leung *et al*., 2002).

To determine the importance of environmental variables in the habitat suitability modeling of Goitered gazelle in the Samelghan plain, the Maxent uses three factors: the Jackknife plot, the percent contribution values of variables, and species response curves to these variables.

According to the Jackknife plot, the variables that were most important in the habitat modeling including: Mediterranean climate class, plateau 3.1 (plateaus with characteristics; medium and low altitudes, medium to high erosion on limestone and sand) and plant type Acantholimon-Astragalus. Also, other environmental variables were also effective in modeling, but the impact of these variables was more noticeable than them. The effect of these variables is such that by removing each of these, there will be significant changes in habitat modeling. According to the Jackknife plot, variables such as residential villages were the variables that had the least effect on the modeling. That is, removing this variable does not make much difference in modeling. The second factor is the percent contribution values table of variables in the habitat modeling. This table shows the participation percentage of each variable in determining the suitable habitat for the studied species at the basin.

As shown in the table, variables such as 0-5% slope class, semi-dense pasture class were among the variables that had the most participation in modeling. Based on this table, it can be determined how much each variable has contributed to creating the final habitat suitability map for Goitered gazelle. After determining the important variables in habitat modeling, based on response curves (third factor) can determine the impact of each variable on the probability of species presence. Based on these curves, it can be determined that increasing or decreasing the distance from the variable, what changes occur in the probability of species presence and habitat suitability.

the results showed that the species showed different responses to different environmental variables.

the slope 0-5% variable; this variable is one of the variables that have the most participation in modeling, based on the species response curve, the probability of species presence decreases with increasing distance from this variable. So, it can be seen that Goitered gazelle prefers areas with a slope of 0-5% compared to other slope classes.

Climatic conditions, in turn, are effective in habitat modeling, according to the Jackknife plot, the Mediterranean climate class is a variable that is more important than other variables. However, based on the response curve of this variable, the probability of the presence of the species changes less, ie with increasing distance from this variable, the probability of presence increases and after a certain distance, there is no change in the probability of presence.

Another important variable in Goitered gazelle modeling is the distance from the semi-dense pasture variable. pastures are very important in the food supply, the Maxent’s results showed that Goitered gazella prefers semi-dense pastures with type Acantholimon-Astragalus vegetation over other variables, as shown in the species response curve to these variables with increasing distance from this variable, the probability of presence much reduced.

The variables obtained from the results of the Jackknife plot and the percent contribution values table for Goitered gazella in the Ashkhneh basin are not necessarily variables that reduce the likelihood of presence. because, if we examine some of the obtained variables based on the response curves, by increasing the distance from it, increases or causes fluctuations in the probability of presence, which can be influenced by other variables or other factors in the habitats such as the existence of competing species, predators and human activities. Variables such as classes of slope 15-30% are among the variables that the probability of species presence will increase with increasing distance from it, so that Goitered gazelle avoids these areas and prefers other slope classes to this variable.

Other variables also have an effect on the probability of species presence, which can be explained by interpreting the response curves to their positive or negative effects on habitat desirability.

The results of the Jackknife plot, the percent contribution values table, provide Maxent a threshold for habitat classification that shows the final suitability map for Goitered gazella in the Ashkhaneh area. The suitability classification map shows that Goitered gazelle’s habitat covers the northern parts of the basin, which has all the suitable conditions for species.

The study showed that approx. 11% of the Samelghan plain is a suitable habitat for Goitered gazelle.

Based on the research, it is suggested that:

- Since the area is not under the protection of the Environment Organization; other species should be identified and examined, if possible, to define as the area as a “No-hunting area”.

## Acknowledgment

The present article is part of the thesis of the master’s degree approved by Yazd University, and the authors thank them for their support. Also, we sincerely thank all people who helped us in this study.

## References

Alatawi, S. A., Gilbert, F., and Reader, T. 2020. Modelling terrestrial reptile species richness, distributions and habitat suitability in Saudi Arabia. Journal of Arid Environments, 178: 104153. https://doi.org/10.1016/j.jaridenv.2020.104153.

Alcala-Canto, Y., Figueroa-Castillo, A. J., Ibarra-Velarde, F., Vera-Montenegro, Y., Cervantes-Valencia, E. M., Salem, M. Z. A., and Cuéllar-Ordaz, A. J. 2018. Development of the first georeferenced map of Rhipicephalus (Boophilus) spp. in Mexico from 1970 to date and prediction of its spatial distribution. Geospat. Health, 13(1): 624. https://doi.org/10.4081/gh.2018.624.

Allouche, O., Steinitz, O., Rotem, D., Rosenfeld, A., and Kadmon, R. 2008. Incorporating distance constraints into species distribution models. Journal of Applied Ecology, 45: 599–609. doi: 10.1111/j.1365-2664.2007.01445.x.

Crooks, R. K., Burdett, L. C., Theobald, M. D., King, R. S., Marco, D. M., Rondinini, C., and Boitani, L. 2017. Quantification of Habitat Fragmentation Reveals Extinction Risk in Terrestrial Mammals. PNAS, 1–6. https://doi.org/10.1073/pnas.1705769114.

Dormann, C. F., Schymanski, S. J., Cabral, J., Chuine, I., Graham, C., Hartig, F., Kearney, M., Morin, X., Römermann, C., Schröder, B., and Singer, A. 2012. Correlation and process in species distribution models: bridging a dichotomy. Journal of Biogeography, 39(12): 2119–2131. https://doi.org/10.1111/j.1365-2699.2011.02659.x.

Elith, J., and Leathwick, J. R. 2009. Species distribution models: ecological explanation and prediction across space and time. Annual Review of Ecology and Systematics, 40: 677–697. https://doi.org/10.1146/annurev.ecolsys.110308.120159.

Elith, J., Phillips, S. J., Hastie, T., Dudik, M., Chee, Y. E., and Yates, C. J. 2011. A statistical explanation of MaxEnt for ecologists. Diversity and Distributions, 17(1): 43–57. https://doi.org/10.1111/j.1472-4642.2010.00725.x.

Evcin, O., Kucuk, O., and Akturk, E. 2019. Habitat suitability model with maximum entropy approach for European roe deer (*Capreolus capreolus*) in the Black Sea Region. Environmental monitoring and assessment, 191, 669. https://doi.org/10.1007/s10661-019-7853-x.

Fand, B. B., Kumar, M., and Kamble, L. A. 2014. Predicting the potential geographic distribution of cotton mealybug Phenacoccus solenopsis in India based on MAXENT ecological niche model. Journal of Environmental Biology, 35: 973–982.

Fielding, A. H., and Bell, J. F. 1997. A review of methods for the assessment of prediction errors in conservation presence/absence models. Environmental Conservation, 24(1): 38–49. https://doi.org/10.1017/S0376892997000088.

Franklin, J., 2013. Species distribution models in conservation biogeography: developments and challenges. Diversity and Distributions, 19(10): 1217–1223. https://doi.org/10.1111/ddi.12125.

Fronczak, L. D., Andersen, E. D., Hanna, E. E., and Cooper, R. T. 2017. Distribution and migration chronology of eastern population sandhill cranes. J Wildlife Manage, 81(6):1021–1032. https://doi.org/10.1002/jwmg.21272.

Ghanbarian, G., Raoufat, R. M., Pourghasemi, R. H., and Safaeian, R. 2019. Habitat suitability mapping of *Artemisia aucheri* boiss based on the GLM model in R. Spatial Modeling in GIS and R for Earth and Environmental Sciences, 213–227. https://doi.org/10.1016/B978-0-12-815226-3.00009-0.

Giovanelli, J. F., Siqueira, F. M., Haddad, B. F. C., and Alexandrino, J. 2010. Modeling a spatially restricted distribution in the Neotropics: how the size of calibration area affects the performance of five presence-only methods. Ecological Modeling, 221(2): 215–224. https://doi.org/10.1016/j.ecolmodel.2009.10.009.

Gormley, A. M., Forsyth, D. M., Griffioen, P., Lindeman, M., Ramsey, D. S. L., Scroggie, M. P., and Woodford, L. 2011. Using presence-only and presence–absence data to estimate the current and potential distributions of established invasive species. Journal of Applied Ecology. 48: 25–34. doi: 10.1111/j.1365-2664.2010.01911.x.

Guilbault, E., Renner, I., Mahony, M., and Beh, E. 2019. Classification of unlabeled observations in Species Distribution Modelling using Point Process Models. bioRxiv, https://doi.org/10.1101/651125.

Guisan, A., and Thuiller, W. 2005. Predicting species distribution: offering more than simple habitat models. Ecology Letters, 8(9): 993–1009. https://doi.org/10.1111/j.1461-0248.2005.00792.x.

Guisan, A., and zimmermann, N. E. 2000. Predictive habitat distribution models in ecology. Ecological Modelling, 135(2-2):147–186. https://doi.org/10.1016/S0304-3800(00)00354-9.

Hernandez, A. P., Graham, H. C., Master, L. L., and Albert, L. D. 2006. The effect of sample size and species characteristics on performance of different species distribution modeling methods. Ecography, 29(5): 773–785. https://doi.org/10.1111/j.0906-7590.2006.04700.x.

Hirzel, A. H., Le Lay, G., Helfer, V., Randin, C., and Guisan, A. 2006. Evaluating the ability of habitat suitability models to predict species presences. Ecological Modelling, 199(2): 142–152. https://doi.org/10.1016/j.ecolmodel.2006.05.017.

Huercha. Song, R., Ma, Y., Hu, Z., Li, Y., Li, M., Wu, L., Li, C., Dao, E., Fan, X., Hao, Y., and Bayin, C. 2020. MaxEnt modeling of *Dermacentor marginatus* (Acari: Ixodidae) distribution in Xinjiang, China. Journal of Medical Entomology, 1–9. https://doi.org/10.1093/jme/tjaa063.

Ito, H., Hayakawa, K., Ooba, M., and Fujii, T. 2020. Analysis of habitat area for endangered species using maxent by urbanization in Chiba, Japan. International Journal of GEOMATE, 18(68): 94–100. https://doi.org/10.21660/2020.68.5721.

Jafari, M., and Thomalski, J. 2016 “The Iranian-German Tappe Rivi Project (TRP), North - Khorasan: Report on the 2016 and 2017 fieldworks”, Archäologische Mitteilungen aus Iran und Turan 48: 7.

Jiménez-Valverde, A. 2012. Insights into the area under the receiver operating characteristic curve (AUC) as a discrimination measure in species distribution modeling. Global Ecology Biogeography, 21(4):498–507. https://doi.org/10.1111/j.1466-8238.2011.00683.x.

Jiménez-Valverde, A., Ortuño, M. V., and Lobo, M. J. 2007. Exploring the distribution of Sterocorax Ortuño, 1990 (*Coleoptera, Carabidae*) species in the Iberian Peninsula. Journal of Biogeography (J. Biogeogr.), 34(8): 1426–1438. https://doi.org/10.1111/j.1365-2699.2007.01702.x.

Kamyo, T., and Asanok, L. 2020. Modeling habitat suitability of *Dipterocarpus alatus* (Dipterocarpaceae) using MaxEnt along the Chao Phraya River in Central Thailand. J. Forest Science and Technology, 16(1): 1–7. https://doi.org/10.1080/21580103.2019.1687108.

Kozak, K. H., Graham, C. H., and Wiens, J. J. 2008. Integrating GIS-based environmental data into evolutionary biology. Trends in Ecology and Evolution, 23(3): 141–148. https://doi.org/10.1016/j.tree.2008.02.001.

Kramer-Schadt, S., Niedballa, J., Pilgrim, D. J., Schroder, B., Lindenborn, J., Reinfelder, V., Stillfried, M., Heckmann, I., Scharf, K. N., Augeri, M. D., Cheyne, M. S., Hearn, J. A., Ross, J., Macdonald, W. D., Mathai, J., Eaton, J., Marshall, J. A., Semiadi, G., Rustam, R., Bernard, H., Alfred, R., Samejima, H., Duckworth, W. J., Breitenmoser-Wuersten, C., Belant, L. J., Hofer, H., and Wilting, A. 2013. The importance of correcting for sampling bias in MaxEnt species distribution models. Diversity and Distributions, (Diversity Distrib.), 19(11): 1366–1379. https://doi.org/10.1111/ddi.12096.

Kuloba, M. B., Gils, V. H., Duren, V. I., Muya, M. S., and Ngene, M. S. 2015. Modeling Cheetah *Acinonyx jubatus* fundamental niche in Kenya. International Journal of Environmental Monitoring and Analysis, 3(5): 317–330. doi:10.11648/j.ijema.20150305.22.

Leung, B., Lodge, M. D., Finnoff, D., Shogren, F. J., Lewis, A. M., and Lamberti, G. 2002. An ounce of prevention or a pound of cure: bioeconomic risk analysis of invasive species. The Royal Society. Biol Sci, 269(1508): 2407–2413. https://doi.org/10.1098/rspb.2002.2179.

Li, J., Fan, G., and He, Y. 2020. Predicting the current and future distribution of three Coptis herbs in China under climate change conditions, using the MaxEnt model and chemical analysis. Science of the Total Environment, 698: 134–141. https://doi.org/10.1016/j.scitotenv.2019.134141.

Lobo, J. M. 2008. More complex distribution models or more representative data?. Biodiversity Informatics. 5:14–19. https://doi.org/10.17161/bi.v5i0.40.

Long, R. P., Zefania, S., Ffrench-Constant, H. R., and Székely, T. 2008. Estimating the population size of an endangered shorebird, the Madagascar plover, using a habitat suitability model. Animal Conservation, 11(2): 118–127. https://doi.org/10.1111/j.1469-1795.2008.00157.x.

Lou, X., Liang, L., Liu, Z., Wang, J., Huang, T., Geng, D., and Chen, B. 2020. Habitat suitability evaluation of the Chinese horseshoe bat (*R. sinicus*) in the Wuling mountain area based on MAXENT modelling. Pol. J. Environ. Stud. 29(1): 1263–1273. DOI: 10.15244/pjoes/102370.

Mashkour, M. 2001. Paleoenvironmental Investigations in the Qazvin Plain (Iran) in Proceedings of the First International Congress of Archaeology of the Near East (ICAANE -May 1998), Rome. 2: 967–982.

Mashkour, M. 2013. Section C. Specialist contributions, Chp. Animal Bones 20.3 Animal exploitation during the Iron Age to Achaemenid, Sasanian and Early Islamic periods along the Gorgan Wall. In Sauer, E., Omrani Rekavandi, H., Wilkinson, T. and Nokandeh J., Persia’s Imperial Power in Late Antiquity: The Great Gorgan Wall and the Frontier Landscapes of Sasanian Iran. British Institute of Persian Studies monograph. British Academy. Oxbow Books. pp:548–580 (bibliography from pp:642-667. ISBN-13:978-1-84217-519-4, ISBN-10:1-84217-519-X.

Mashkour, M., Fontugne, M. And Hatté, C. 1999. Investigations on the Evolution of Subsistence Economy in the Qazvin Plain (Iran) from the Neolithic to the Iron Age. Antiquity, 73(279): 65–76. doi:10.1017/S0003598X00087846.

Merow, C., Smith, J. M., and Silander, A. J. 2013. A practical guide to MaxEnt for modeling species’ distributions: what it does, and why inputs and settings matter. Ecography, 36(10): 1058–1069. https://doi.org/10.1111/j.1600-0587.2013.07872.x.

Merow, C., Smith, J. M., Edwards, C. T., Guisan, A., Mcmahon, M. S., Normand, S., Thuiller, W., Wüest, O. R., Zimmermann, E. N., and Elith, J. 2014. What do we gain from simplicity versus complexity in species distribution models?. Ecography, 37(12):1267–1281. https://doi.org/10.1111/ecog.00845.

Mi, C., Falk, H., and Guo, Y. 2016. Climate envelope predictions indicate an enlarged suitable wintering distribution for Great Bustards (Otis tarda dybowskii) in China for the 21st century. Peer J, 4:e1630. https://doi.org/10.7717/peerj.1630.

Moreno, R. Zamora, R., Molina, R. J., Vasquez, A., and Herrera, A. M. 2011. Predictive modeling of microhabitats for endemic birds in South Chilean temperate forests using Maximum entropy (Maxent). Ecological Informatics, 6(6): 364–370. https://doi.org/10.1016/j.ecoinf.2011.07.003.

Mousazade, M., Ghorbanian, G., Pourghasemi, R. H., Safaeian, R., and Cerda, A. 2019. maxent data mining technique and its comparison with a bivariate statistical model for predicting the potential distribution of *Astragalus Fasciculifolius* Boiss. in Fars, Iran. Sustainability, 11(12). 3452, https://doi.org/10.3390/su11123452.

Muscarella, R., Galante, J. P., Soley-Guardia, M., Boria, A. R., Kass, M. J., Uriarte, M., and Anderson. P. R. 2014. ENMeval: An R package for conducting spatially independent evaluations and estimating optimal model complexity for M AXENT ecological niche models. Methods in Ecology and Evolution, 5(11): 1198–1205. https://doi.org/10.1111/2041-210X.12261.

Naimi, B., Hamm, A. S. N., Groen, A. T., Skidmore, K. A., and Toxopeus, G. A. 2014. Where is positional uncertainty a problem for species distribution modelling?. Ecography, 37(2): 191–203. https://doi.org/10.1111/j.1600-0587.2013.00205.x.

Naqibzadeh A, Sarhangzadeh J, Sayedi N. 2021. Habitat desirability modeling of Goitered Gazelle (Gazella subgutturosa) by Ecological Niche Factor Analysis in the Bidouyeh Protected Area, Iran. Journal of wildlife and Biodiverity, 5(4):15–27. 10.22120/JWB.2021.528662.1223.

Neer, V. W. 2017. Archaeozoology in Sub-Saharan Africa. Field Manual for African Archaeology. Smith, L, A. Cornelissen, E. Gosselain, P,O. MacEachern, S (Edts). Royal Museum for Central Africa, Tervuren 2017. Chapter 5: 210–213.

Ortega-Huerta, A. M., and Peterson, T. A. 2008. Modeling ecological niches and predicting geographic distributions: a test of six presence-only methods. Revista Mexicana de Biodiversidad, 79:205–216. http://www.redalyc.org/articulo.oa?id=42579117.

Pearce, J. L., and Boyce, M. S. 2006. Modelling distribution and abundance with presence-only data. J. Appl. Ecol, 43(3): 405–412. https://doi.org/10.1111/j.1365-2664.2005.01112.x.

Pearce, J., and Ferrier, S. 2000. Evaluating the predictive performance of habitat models developed using logistic regression. Ecol Model, 133(3): 225–245. https://doi.org/10.1016/S0304-3800(00)00322-7.

Pearson, G. R., Raxworthy, J. C., Nakamura, M., and Peterson, T. A. 2007. Predicting species distributions from small numbers of occurrence records: a test case using cryptic geckos in Madagascar. J. Biogeography, 34(1): 102–117. https://doi.org/10.1111/j.1365-2699.2006.01594.x.

Pédarros, É., Coetzee, T., Fritz, H., and Guerbois, C. 2020. Rallying citizen knowledge to assess wildlife occurrence and habitat suitability in anthropogenic landscapes. Biological Conservation, 242, 108407. https://doi.org/10.1016/j.biocon.2020.108407.

Pédarros, E., Goetzee, T., Fritz, H., and Guerbois, C. 2020. Rallying citizen knowledge to assess wildlife occurrence and habitat suitability in anthropogenic landscapes. Biological Conservation, 242: 108407. https://doi.org/10.1016/j.biocon.2020.108407.

Peterson, A. T., Soberon, J., Pearson, R. G., Anderson, R. P., Martinez-Meyer, E., Nakamura, M. and Araujo, M. B. 2011. Ecological niches and geographic distributions. Princeton University Press, Princeton, NJ.

Phillipes, S, J,. 2012. A brief tutorial on Maxent, versions. 3.3.3. Available online: http://www.cs.princeton.edu/~schapire/maxent/.

Phillips, J, S., Anderson, P, R., Dudík, M., Schapire, E. R., and Blair, E. M. 2017. Openingthe black box: An open source release of Maxent. Ecography, 40. https://doi.org/10.1111/ecog.03049.

Phillips, J. S., and Dudik, M. 2008. Modeling of species distributions with MaxEnt: new extensions and a comprehensive evaluation. Ecography, 31(2): 161–175. https://doi.org/10.1111/j.0906-7590.2008.5203.x.

Phillips, J. S., and Dudik, M. 2008. Modeling of species distributions with MaxEnt: new extensions and a comprehensive evaluation. Ecography, 31(2) 161–175. doi: 10.1111/j.2007.0906-7590.05203.x.

Phillips, J. S., Anderson, P. R., Schapire, E. R. 2006. Maximum entropy modeling of species geographic distributions. Ecological Modelling, 190(3-4): 231–259. https://doi.org/10.1016/j.ecolmodel.2005.03.026.

Phillips, L. S., Dudík, M., and Schapire, R. 2004. A Maximum Entropy Approach to Species Distribution Modeling. Proceedings of the 21th International Conference on Machine Learning. Banff, Canada. https://doi.org/10.1145/1015330.1015412.

Phillips, S, J., Dudik, M., Schapire, R, E. 2004. A maximum entropy approach to species distribution modeling. In: Proceed of the 21st Int. conf. on Machine Learning, AcM Press, New York. pp:655–662. https://doi.org/10.1145/1015330.1015412.

Phillips, S., Anderson, R. R., and Schapire, R. 2006. Maximum entropy modeling of species geographic distributions. Ecol Model. 190(3-4): 231–259. https://doi.org/10.1016/j.ecolmodel.2005.03.026.

Preau, C., Trochet, A., Bertrand, R., and Isselin-Nondedeu, F. 2018. Modeling potential distributions of three European amphibian species comparing Enfa and Maxent. Herpetological Conservation and Biology, 13(1): 91–104.

Qin, A., Jin, K., Batsaikhan, E. M., Nyamjav, J., Li, G., Li, J., Xue, Y., Sun, G., Wu, L., Indree, T., Shi, Z., and Xiao, W. 2020. Predicting the current and future suitable habitats of the main dietary plants of the Gobi Bear using MaxEnt modeling. Global Ecology and Conservation, 22: e01032. https://doi.org/10.1016/j.gecco.2020.e01032.

Qin, A., Liu, B,. Guo, Q., Bussmann, R.W., Ma, F., Jian, Z., Xu, G., and Pei, S. 2017. Maxent modeling for predicting impacts of climate change on the potential distribution of Thuja sutchuenensis Franch., an extremely endangered conifer from southwestern China. Glob. Ecol. Conserv, 10: 139–146. https://doi.org/10.1016/j.gecco.2017.02.004.

Radosavljevic, A., and Anderson, P. R. 2014. Making better MAXENT models of species distributions: complexity, overfitting and evaluation. Journal of Biogeography (J. Biogeogr.), 41(4): 629–643. https://doi.org/10.1111/jbi.12227.

Renner, W. I., Elith, J., Baddeley, A., Fithian, W., Hastie, T., Phillips, J. S., Popovic, G., and Warton, D. I. 2015. Point process models for presence-only analysis. Methods in Ecology and Evolution, 6(4): 366–379. https://doi.org/10.1111/2041-210X.12352.

Ruete, A., and Leynaud, C. G. 2015. Goal-oriented evaluation of species distribution models’ accuracy and precision: True skill statistic profile and uncertainty maps. PeerJ PrePrints. https://dx.doi.org/10.7287/peerj.preprints.1208v1.

Saupe, E. E., Qiao, H., Hendricks, R. J., Portell, W. R., Hunter, J. S., Soberón, J., and Lieberman, S. B. 2015. Nichebreadthand geographic Range size as determinants of species survival on geological time scales. Global Ecology and Biogeography, 24(10): 1159–1169. https://doi.org/10.1111/geb.12333.

Signorini, M., R. Cassini, M. Drigo, A. Frangipane di Regalbono, M. Pietrobelli, F. Montarsi, and A. S. Stensgaard. 2014. Ecological niche model of Phlebotomus perniciosus, the main vector of canine leishmaniasis in north-eastern Italy. Geospat. Health. 9(1): 193–201. https://doi.org/10.4081/gh.2014.16.

Suleman, S., Khan, A. W., Anjum, M. K., Shehzad, W., and Hashemi, M. G. S. 2020. HABITAT suitability index (HSI) model of Punjab urial (*Ovis vegnei punjabiensis*) in Pakistan. The J. Anim. Plant Sci, 30(1).

Thomalsky, J. 2016. Tappeh Rivi, Iran: Die iranisch-deutschen arbeiten des Jahres 2016. e - Forschungsberichte des dai, 3: 2–11. http://nbn-resolving.de/urn:nbn:de:0048-dai-edaif.2016-3-1543.(Germany)

Thomas, L, Buckland, S. T., Rexstad, E. A., Laake, J. L,. Strindberg, S., Hedley, S. L., Bishop, J. R, B., Marques, T. A., and Burnham, K. P. 2010. Distance software: design and analysis of distance sampling surveys for estimating population size. J Applied Ecology, 47: 5–14. https://doi.org/10.1111/j.1365-2664.2009.01737.x.

Toor, L. M., Jaberg, C. and Safi, K,. 2011. Integrating sex-specific habitat use for conservation using habitat suitability models. Animal Conservation, 14(5): 512–520. https://doi.org/10.1111/j.1469-1795.2011.00454.x.

Tourne, M. C. D., Ballester, R. V. M., James, A. M. P., Martorano, G. L., Guedes, C. M., and Thomas, E. 2019. Strategies to optimize modeling habitat suitability of *Bertholletia excelsa* in the Pan-Amazonia. Ecology and Evolution, 9(22): 12623–12638. https://doi.org/10.1002/ece3.5726.

Traill, W. L., and Bigalke, C. R. 2007. A presence-only habitat suitability model for large grazing African ungulates and its utility for wildlife management. Afr. J. Ecol., 45: 347–354.

Van Strien, J. A., Van Swaay, M. A. C., and Termaat, T. 2013. Opportunistic citizen science data of animal species produce reliable estimates of distribution trends if analysed with occupancy models. J. Applied Ecology, 50(6): 1450–1458. https://doi.org/10.1111/1365-2664.12158.

Waltert, M., Meyer, B., Shanyangi, M. W., Balozi, J. J., Kitwara, O., Qolli, S., Krischke, H., and Muehlenberg, M. 2008. Foot surveys of large mammals in woodlands of western Tanzania. Journal of Wildlife Management, 72(3): 603–610. https://doi.org/10.2193/2006-456.

Wan, J. Z., Wang, C. J., Yu, J. H., Nie, S. M., Han, S. J., Liu, J. Z., Zu, Y. G., and Wang, Q. G., 2016. Developing conservation strategies for Pinus koraiensis and Eleutherococcus senticosus by using model-based geographic distributions. Journal of Forestry Research, 27: 389–400. https://doi.org/10.1007/s11676-015-0170-5.

Wan, J., Wang, C., and Yu, F. 2019. Effects of occurrence record number, environmental variable number, and spatial scales on MaxEnt distribution modelling for invasive plants. Biologia, 74: 757–766. https://doi.org/10.2478/s11756-019-00215-0.

Wang, J., Liu, H., Li, Y., and Zhang, H. 2019. Habitat quality of overwintering red-crowned cranes based on ecological niche modeling. Arabian Journal of Geosciences, 12:750. https://doi.org/10.1007/s12517-019-4932-9.

Warren, L. D., Glor, E. R, and Turelli, M. 2008. Environmental niche equivalency versus conservatism: quantitative approaches to niche evolution. Evolution, 62(11):2868–2883. https://doi.org/10.1111/j.1558-5646.2008.00482.x.

Warton, I. D,. and Shepherd, C. L. 2010. Poisson point process models solve the “pseudo-absence problem” for presence-only data in ecology. The Annals of Applied Statistics, 4(3): 1383–1402. doi:10.1214/10-AOAS331.

Xu, L. Z., Peng, H. H., and Peng, Z. S. 2015. The development and evaluation of species distribution models. Acta Ecologica Sinica, 35:557–567.

Yackulic, B. C., Chandler, R., Zipkin, F. E., Royle, A. J., Nichols, D. J., Grant, C. H. E., and Veran, S. 2013. Presence-only modelling using MAXENT: when can we trust the inferences?. Methods in Ecology and Evolution, 4(3): 236–243. https://doi.org/10.1111/2041-210x.12004.

Yost, A. C,. Petersen, S. L,. Gregg, M., and Miller, R. 2008. Predictive modeling and mapping sage grouse (*Centrocercus urophasianus*) nesting habitat using maximum entropy and a long-term dataset from southern Oregon. Ecological Information. 3(6): 375–386. https://doi.org/10.1016/j.ecoinf.2008.08.004.

Yuan, H. S., Wei, Y. L., & Wang, X. G. (2015). Maxent modelling for predicting the potential distribution of Sanghuang, an important group of medicinal fungi in China. Fungal Ecology, 17, 140–145. https://doi.org/10.1016/j.funeco.2015.06.001.

Zhang, J., Jiang, F., Li, G., Qin, W. Li, S., Gao, H., Cai, Z., Lin, G., and Zhang, T. 2019b. Maxent modeling for predicting the spatial distribution of three raptors in the Sanjiangyuan National Park, China. Ecology and Evolution, 9(11): 6643–6654. https://doi.org/10.1002/ece3.5243.

Zhang, K., Yao, L., Meng, J., and Tao, J. 2018. Maxent modeling for predicting the potential geographical distribution of two peony species under climate change. Science of the Total Environment, 634, 1326–1334. https://doi.org/10.1016/j.scitotenv.2018.04.112.

Zhang, Z. X., Capinha, C., Weterings, R., McLay, C. L., Xi, D., Lu, H. J., and Yu, L. Y., 2019a. Ensemble forecasting of the global potential distribution of the invasive Chinese mitten crab, Eriocheir sinensis. Hydrobiologia, 826: 367–377. https://doi.org/10.1007/s10750-018-3749-y.

